# Angiotensin II type 1 receptor activation facilitates pain hypersensitivity via dorsal horn pericyte mediated vasoconstriction

**DOI:** 10.1101/2023.12.12.571346

**Authors:** Lydia D. Hardowar, Matt Sheavyn, Philip G. McTernan, Dave O. Bates, Richard P. Hulse

**Affiliations:** School of Science and Technology, Nottingham Trent University, UK; Division of Cancer and Stem Cells, School of Medicine, Centre for Cancer Science, Biodiscovery Institute, University of Nottingham, Nottingham, NG2 7UH, UK; Centre of Membrane and Protein and Receptors (COMPARE), University of Birmingham and University of Nottingham, Midlands, UK

**Keywords:** pain, pericyte, spinal cord, endothelial, angiotensin II

## Abstract

Vascular disturbance is a key factor in the development of neurological disease, with reduced integrity of the capillary network in the dorsal horn implicated in activation of nociceptive neural circuits and induction of pain hypersensitivity. Pericytes regulate capillary health and tone, with pericyte dysfunction in cerebral tissue associated with neurodegenerative disorders. Our work demonstrates that nociceptive processing is influenced by angiotensin II type 1 (AT1) receptor mediated pericyte contractility in the dorsal horn. Intravital imaging of the mouse spinal cord demonstrated angiotensin II induced cessation of spinal cord capillary perfusion. Intrathecal administration of angiotensin II induced pericyte contractility and narrowing of capillary diameter, which was accompanied by mechanical allodynia and heat hyperalgesia. Angiotensin II mediated pericyte activation and reduction of spinal cord blood flow, was prevented by inhibition of AT1 receptor via losartan treatment. In addition, losartan either systemically or intrathecally administered, prevented angiotensin II induced pain in male and female adult mice. This was associated with protection of the dorsal horn capillary endothelium, with intrathecal co-treatment with losartan preventing loss of CD31 immunoreactivity in the dorsal horn following administration of angiotensin II. This investigation demonstrates that AT1 mediated pericyte regulation of the dorsal horn capillary network, is fundamental in modulating nociceptive processing and perception of pain. Here we identify a novel cellular and mechanistic target for the development of new analgesic.

## Introduction

Chronic pain is a debilitating affliction impacting large proportions of the global population [1,9], with presentation typified as exaggerated perception of pain to differing sensory modalities and accompanying ongoing pain experiences. Furthermore, this is accompanied by an aversive and negative emotive nature of the chronic pain experience, which heavily impacts upon an individual’s quality of life. The transition from acute protective pain to neuropathic pain is classified by damage caused to the somatosensory nervous system, typically by either disease or treatment. This has inherently led to neuronal damage as a primary target for investigations into chronic pain. However, it is widely regarded that in times of chronic pain following traumatic injury or due to inflammatory intervention, alterations in the blood spinal cord barrier permeability facilitates the infiltration of the circulatory inflammatory cells (Beggs *et al*., 2010). Additionally, the somatosensory nervous system is one of the most metabolically active tissues, which is reliant upon a continual blood supply of oxygen and glucose to support the extensive anatomical and functional processes that are essential to maintain nociception and somatosensation. As a consequence, damage to this vascular system can be devastating, with degeneration of the endothelium heavily associated with neurodegenerative disease(Nortley *et al*., 2019). Our recent work has highlighted in mouse models that by inducing damage to the endothelium in the dorsal horn (i.e loss of blood vessels), directly causes pain hypersensitivity. Additionally, recent evidence highlights dysfunction in the adjoining mural cells that support the endothelial cells in relation to function and integrity, are implicated in neurodegenerative disease(Nortley *et al*., 2019). For example, pericyte mediated vasocontractility underlies reduced neural tissue perfusion (Li *et al*., 2017; Berthiaume *et al*., 2018) in Alzheimers and stroke (Hall *et al*., 2014*a*; Li *et al*., 2017; Nortley *et al*., 2019). Importantly, there is a strong association between the prevalence of chronic pain states and incidence of hypertension(Ponirakis *et al*., 2021). The renin angiotensin system (RAS) is widely recognised as the principle system that governs vascular tone during hypertension, with angiotensin receptor type 1 (AT1) a therapeutic target. Pericytes are in close apposition to the endothelium governing the function and integrity, and fundamentally in modulating capillary tone, with angiotensin II (AngII) a potent vasoconstrictor (Nadal *et al*., 2002; Kamouchi *et al*., 2004; Hirunpattarasilp *et al*., 2021), is a putative driver of pericyte contractility. AngII and the vasoconstrictor modulating receptor angiotensin II type 1 (AT1) are expressed in spinal cord (Ahmad *et al*., 2003; Wosik *et al*., 2007) implying a potential for modulation nociceptive processing and pain development.

Our investigations aim to explore the functional effects of AngII on spinal blood flow during chronic pain development, determining the impact of AT1 mediated pericyte-induced vasocontractility. Here we demonstrate that angiotensin II mediated pericyte contractility is AT1 dependent and curtails dorsal horn capillary perfusion to induce chronic pain.

## Materials and Methods

### Ethical Approval and Animals Used

All experiments were carried out in accordance with the United Kingdom’s Scientific Procedures Act 1986 and were approved by the Nottingham Trent University and University of Nottingham Animal Welfare and Ethical Review Boards (AWERB). Adult C57/BL6J mice (6-8 weeks old) male and female mice were used (male n=39, female n=18). Adult C57/BL6J male mice (8 weeks old n=20) were used for Laser speckle contrast imaging analysis and (8 weeks old n=20) were used for intravital confocal imaging. All mice were housed with environmental enrichment under 12:12 hour light dark cycles. All mice had ad libitum access to water and standard diet chow (Rodent 2018 Envigo Global Certified Diet: USA). C57/BL6J male and female pups were used at postnatal 7-14 day for primary spinal cord pericyte cultures *in vitro*.

### Drug Preparation

Angiotensin II (AngII), Losartan potassium and PD123319 ditrifluoroacetate (Tocris Bioscience; UK) were prepared in sterile distilled phosphate buffer saline (PBS). Final concentrations were prepared and administered to mice as outlined vehicle groups of age-matched mice received PBS treatments. AngII was administered intrathecally (i.t.) at 100nM. Depending on the study, randomised cohorts mice were selected for intraperitoneal (20mg/kg; i.p.) or i.t. delivery of losartan (10µM) or i.p. PD123319 (10µM) alongside i.t. AngII (details of animal use summarised in Table 1). Drug concentration administered to rodents were based upon previous studies presented *in vitro* (Shepherd *et al*., 2018*a*, 2018*b*).

**Table 1.** Details of animals within the AngII-induced pain model Details of drug name (PBS, AngII, Losartan, PD123319) and drug delivery route (i.t. or i.p.) are stated for each experimental group. Known drug concentrations for single drug dose or combined mixture are represented with drug name alongside animal group information (animal number and sex).

| Experimental group<br>drugs with delivery<br>route(s) | Drug concentration | Animal number | Sex |
| --- | --- | --- | --- |
| i.t. veh | 0.01M (PBS) | 15 | 6 female<br>9 male |
| i.t. AngII | 100nM | 15 | 6 female<br>9 male |
| i.t. AngII + i.p. vehicle | 100nM (AngII) +<br>0.01M (veh) | 6 | male |
| i.t. AngII + i.p.<br>PD123319 | 100nM (AngII) +<br>10mg/kg (PD123319) | 3 | male |
| i.t. AngII + i.p.<br>Losartan | 100nM (AngII) +<br>20mg/kg (Losartan) | 6 | male |
| i.t. AngII + i.t. Losartan | 100nM (AngII) + 10µM<br>(Losartan) | 12 | 6 female<br>6 male |

### Intrathecal Administration

Mice were anaesthetised using inhalation anaesthetic (∼2% isoflurane O_2_). Prior to i.t. injection, animals were confirmed to be areflexive and were regularly monitored throughout. An i.t. injection was introduced between intervertebral spaces between lumbar segments 5 and 6. 20µL volume of drug was administered using an insulin needle (BD Micro-Fine 30G). Animals were monitored and allowed to recover. All animals did not present any compromised motor function. During these studies no animal presented alterations in behavioural activity post injection. Studies involving additional i.p. injections for experimental treatments including vehicle (PBS), losartan or PD123319, animals were immediately injected post i.t. injection using a 27 gauge needle insulin syringe. Following injections, nociceptive behavioural testing was performed within 1, 5 and 24 hours.

### Intravital Spinal Cord Imaging

Intravital imaging was performed as previously described (Lobo *et al*., 2023). C57BL6 adult male mice (∼25-30g) were anesthetised via i.p. injection of Ketamine (50mg/kg) and Medetomidine hydrochloride (0.5mg/kg). Rodents were regularly monitored, that they were areflexive and body temperature (∼37°C) was maintained using a feedback control heat blanket. A laminectomy was performed between T12-L1 to allow access to lumbar regions L3-L5 and an in house window chamber was attached allowing positioning of a glass coverslip (diameter = 5mm, thickness = #0, Warner Instruments) to be inserted over the region of interest. The coverslip was secured using silicone elastomer. Capillaries and endothelium were identified using fluorescent tracers introduced via intravenous (tail vein). AlexaFluor 555 conjugated wheat germ agglutinin, (WGA555, 4mg/kg, equating to a volume of 100uL in sterile PBS per mouse, ThermoFisher Scientific, Inc cat no. W32464) and sodium fluorescein (Sigma-Aldrich, cat no.1038870050, 100mg/ml) were administered in sterile PBS via intraperitoneal injection. 15 minutes post injection the mouse was positioned on the microscope stage. The spinal cord was imaged via the window chamber using an inverted confocal microscope (SP8, Leica Microsystems) with a bespoke physiological stage (Scientifica) to hold manipulators. Individual vessels were identified as containing WGA555 labelled endothelium, diameter less than 20μm and identified vessel segment had no branch points along the length to be used for analysis.

### Nociceptive Behaviour

Rodents were habituated to experimenter handling and experimental environment prior to the onset of nociceptive behavioural testing. Nociceptive behavioural assays, Von Frey and Hargreaves tests, were performed to evaluate nociceptive behavioural phenotypes and as previously described (Vencappa *et al*., 2015; Bestall *et al*., 2018; Ved *et al*., 2018*b*).

### Mechanical nociceptive behavioural assay

For von Frey testing, the mice were placed within clear modular holder Perspex cages (Bioseb; USA). The enclosures were placed on top of mesh flooring to enable access for application of von Frey monofilaments (Bioseb; USA) to the medial paw plantar surface of the mouse hindpaw. A range of von Frey monofilaments with increasing force (0.16-2g) was applied to the hindpaw. Each Von Frey was applied to a hindpaw for a minimum interval of 5 minutes. Each force of filament was repeated five times on each paw. The five repeat recordings from each set of increasing filament force (ranging between 0.16-2g) were plotted as a percentage of 0-100% withdrawal response. The data from both left and right paws were merged for each mouse to generate a withdrawal response curve and 50% mechanical nociceptive withdrawal thresholds determined.

### Heat nociceptive behavioural assay

For Hargreaves testing(Hargreaves *et al*., 1988), the mice were placed within clear Perspex enclosures on top of a transparent framed glass pane. A heat source was placed beneath the glass panel positioned under the hindpaw and the latency was recorded upon hindpaw withdrawal.

### Laser speckle imaging blood flow analysis

Animals were anesthetised using inhalation anaesthetic (∼2% isoflurane O_2_) and maintained areflexive for duration of the procedure. Rodent body temperature was maintained at ∼37°C using a heat blanket during the procedure with control feedback delivered via rectal probe to maintain constant temperature. Once anaesthetised, the skin along the dorsal surface of the mouse was shaved. Using a scalpel, an incision was made in the skin along the midline. Muscle, and connective tissue surrounding the spinal vertebra were removed. Laminectomy was performed to reveal the lumbar spinal cord. Placed underneath the moorFLPI-2 instrument, the exposed superficial dorsal longitudinal region of the spinal cord was selected for view. Laser speckle contrast imaging is able to produce an interference pattern between multiple scatter optics fields when light is emitted to a surface of imperfections i.e. moving red blood cells (Dainty, 1977; Wang *et al*., 2007). The selected of view at the lumbar region between L1 to L7 represented the region of interest (ROI). The ROIs were previewed to allow laser speckle camera image optimisation through change in gain and focus before image capture. Once set, images of the spinal cord were taken at 1, 3, 5 and 10-minutes post topical application of experimental drug (Montezano *et al*., 2014). Analysis of images occurred via the moorFLPI-2research software with the generated flux mean of each ROI plotted as percentage change from baseline.

### Immunofluorescence

Mice were terminally anaesthetised with sodium pentobarbital (200mg/kg; Dolethal) and cardiac perfused with PBS and then subsequently with 4% paraformaldehyde. Lumbar spinal cord was extracted and further submerged in 4% PFA for 30 minutes. Spinal cord tissues were washed in PBS and transferred to 30% sucrose solution incubation overnight at 4°C for cryoprotection. After incubation, spinal cord tissues were embedded in optimal cutting temperature compound (VWR International, UK) and stored at −80°C or processed. Embedded SC were sectioned transversely at 50µm thickness using a Leica cryostat CM1860 UV (UK). Serial sectioning was performed to determine tissue change at different measured depths of the lumbar regions of the sections taken, sequential cutting of 50µm tissue sections were mounted onto a SuperFrost Plus Microscope Slide (VWR International; UK). Slides were washed 3 times with PBS for 5 minutes. Tissue sections were permeabilised for 5 minutes in permeabilising solution (0.2% Triton X-100/ PBS) and incubated in blocking solution (5% Bovine serum albumin/0.2% Triton X-100 solution) for 1 hour at room temperature. Following this, primary antibodies were appropriately diluted in blocking solution and left to incubate for 72 hours at 4°C. A no primary control was included as a negative control. Slides were washed 3 times with PBS for 5 minutes. All slides were incubated in secondary antibodies at the appropriate dilution in permeabilising solution at room temperature for 2 hours. Secondary antibodies were removed, and slides were washed twice with PBS for 5 minutes. DAPI (Merck; UK 28718-90-3) was included as a final wash stage. Slides were mounted with coverslips using Vectashield anti-fade mounting media (H1000, Vector laboratories; UK) and stored at 4°C, light protected, until imaged.

### Primary mouse cell isolation

All plates required for cell culture were coated with 100µg/mL with collagen rat tail type I (20nM acetic acid/distilled water) overnight before seeded with cells (Sigma-Aldrich; UK, ThermoFisher Scientific; USA). Spinal cords were collected from postnatal 7–14 day old mice and placed in Hanks balance buffer solution (Gibco; UK) on ice. Tissues were enzymatically dissociated with 0.125% collagenase (Sigma-Aldrich; UK) at 37°C for 15 minutes. Tissues were further dissociated with mechanical force via trituration 10 times. This was repeated 3-4 times. A P200 pipette was used with the corresponding pipette tip to transfer only the cell suspension and placed onto a 20% BSA cushion (40% BSA solution [Sigma-Aldrich; UK], preheated at 37°C) and centrifuged at 3000g for 10 minutes. The supernatant, including tissue debris and myelin layer were discarded, and the pellets re-suspended in preheated pericyte specific culture (PSC) media. (Pericyte specific media containing: 50% DMEM [Gibco; UK], 50% Hams F12 medium [Gibco; UK], 20% fetal bovine serum [Gibco; UK], 1% pericyte growth supplement [Promocell; USA] and 1x penicillin/streptomycin [Gibco; UK]). Cell suspensions were passaged out onto coated 6, 24 or 96 well plates. The cells were incubated within CO_2_ incubator at 37°C overnight before a half media change was performed. The cells were required to reach 80% confluency before cell splitting proceeded. Cells were maintained by full PSC media change every 2 days to replenish nutrient content of PSC media.

### Primary Cell Culture

Once cells had reached a minimal cell confluency of 80%, cells were prepared for experimental use. PSC media in each well were discarded and each well washed with sterile PBS 3 times. 0.05% Trypsin (Sigma-Aldrich; UK) was incubated at 37°C for 5 minutes for cell detachment. Trypsin was neutralised with complete media and centrifuged at 1000rpm for 5 minutes (Eppendorf Centrifuge 5702, U.K.). Post centrifugation, cell pellets were resuspended in PSC media and seeded at 50,000 cells/mL cell density in 6 well plate, 20,000 cells/mL in 24 well plates and 2000 cells/mL in 96 well plates. The LUNA-II automated cell counter (Logo Biosystems; France) was used to perform automated cell count on cell suspensions containing half dilution of Trypan blue (0.2%, ThermoFisher Scientific; USA). Vitality of the cultured cells were assessed at this stage through the live to dead cell count ratio as determined by the LUNA-II automated cell counter.

### Confocal Imaging

All tissue slides and cell coverslips were imaged using a Leica SP5 confocal microscope (Germany). Slides and plates were imaged on a XYZ plane with 25 z-stacks per image. Images were formatted to be the resulting size of 1024×1024 pixels per image, imaged at 400Hz.

### Calcium activity assay

To determine AngII induced activity, Fluo-4-Direct assay (ThermoFisher Scientific; USA) was utilised to measure intracellular calcium concentration in isolated mouse spinal cord pericytes plated in a 96 well plate. Fluo-4 Direct was used as outlined by the manufacturer utilising a 2x Fluo-4 Direct calcium reagent loading. Within the 96 well plates, 2x Fluo-4 Direct calcium reagent loading solution was added to each well and left to incubate for 60-minutes at 37°C. Prior to each drug treatment, readings were taken of each well, which would later be utilised for baseline readings set at T=0. All plate readings were performed using the Tecan plate reader Infinite 200Pro (Life sciences; UK). The settings of the plate reader were set to 37°C, the wavelength settings were selected at 488nm excitation and 516 emission, with the gain set to 180. Experimental drugs (PBS, AngII or Losartan) were added to each well and immediately measured post treatment. The fluorescence response per well was measured at 10 second intervals over a 300 second time period post drug application. The data obtained was represented as fluorescent change over-time (F/F_0_) and area under the curve.

### Immunocytochemistry

Primary pericyte cells were required to be at passage 2-4 for immunocytochemistry. This increase in passage number allowed a reduction in number of endothelial cell and glia cell, as suggested in previous papers (Tigges *et al*., 2013). Immunofluorescence was performed within 24 well plates containing rounded 13mm glass coverslips. Glass coverslips were sterilised with 70% industrial methylated spirit for 5 minutes and washed in PBS before being placed within the wells of a 24 well plate. Media within each well was discarded and washed 3 times with PBS. 1% PFA was used to fix the cells for 30 minutes at room temperature. PFA was discarded from the wells and cells were washed thrice as previously stated. Cells were permeabilised for 5 minutes in PBS 0.2% Triton x-100 and blocked in blocking solution (5%BSA/ 0.2% Triton x-100/PBS) for 1 hour at 4°C. Primary antibodies (Table 2) were appropriately diluted in blocking solution and incubated for 1 hour at room temperature. Primary antibodies were removed, and wells were washed within PBS 3 times for 5 minutes. Wells were incubated in secondary antibodies (0.2% Triton x-100/PBS) at room temperature for 1 hour, light protected. Secondary antibodies were removed, and wells were washed twice with PBS for 5 minutes. A final 5 minute incubation occurs with DAPI wash. Coverslips were carefully removed and mounted on a SuperFrost microscope slide with Vectashield anti-fade mounting media. Slides were stored at 4°C until imaged.

**Table 2.** Antibodies used for cellular and tissue immunofluorescent staining and western blotting. The target antibody, antibody species raised, antibody mix dilution factor and supplier are provided for all primary and secondary antibodies utilised within immunofluorescent staining, Additionally the antibody mix, composition of antibody mix, dilution factor of mix and supplier of reagents are provided for primary and secondary antibodies utilised for western blot protein labelling. (AT1: Angiotensin II type 1, CD31: platelet endothelial cell adhesion molecule, COLIV: collagen IV, IgG: Immunoglobin G, NeuN: neuronal nuclear protein, NG2: chondroitin sulfate proteoglycan, PBS-[T]: phosphate buffer saline-[Tween], SMA: smooth muscle actin).

| Target | Species | Dilution Factor | Supplier |
| --- | --- | --- | --- |

| Primary Antibody |  |  |  |
| --- | --- | --- | --- |
| Anti-CD31 | Goat | 1:200 | R&D systems; UK |
| Anti-NG2 | Rabbit | 1:200 | Millipore; UK |
| Anti-AT1 | Mouse | 1:100 | Santa Cruz; USA |
| Anti-COLIV | Rabbit | 1:500 | Abcam; UK |
| Anti-NeuN | Guinea-pig | 1:500 | Synaptic Systems; USA |
| Anti-C-fos | Rabbit | 1:100 | Santa Cruz; USA |
| Anti-rabbit | Biotinylated | 1:1000 | R&D systems; UK |
| Anti-SMA Cy3 | Conjugated | 1:500 | Abcam; UK |
| Secondary antibodies |  |  |  |
| Anti-goat | Donkey IgG (H+L) | 1:1000 | Invitrogen; USA<br>Alexa Fluor 488 |
| Anti-goat | Donkey IgG (H+L) | 1:1000 | Invitrogen; USA<br>Alexa Fluor 555 |
| Anti-goat | Donkey IgG (H+L) | 1:1000 | Invitrogen; USA<br>Alexa Fluor 647 |
| Anti-rabbit | Streptavidin | 1:1000 | Invitrogen; USA<br>Alexa Fluor 555<br>conjugated |
| Anti-rabbit | Donkey IgG (H+L) | 1:1000 | Invitrogen; USA<br>Alexa Fluor 647 |
| Anti-mouse | Donkey IgG (H+L) | 1:1000 | Invitrogen; USA<br>Alexa Fluor 488 |
| Anti-mouse | Donkey IgG (H+L) | 1:1000 | Invitrogen; USA<br>Alexa Fluor 555 |
| Anti-guinea pig | Goat IgG (H+L) | 1:1000 | Invitrogen; USA<br>Alexa Fluor 488 |
| Western blot antibody mix |  |  |  |
| Antibody mix | Composition | Dilution Factor | Supplier |
| Primary | Anti- AT1 mouse<br>Anti- β-actin rabbit<br>in 5% dried<br>milk/PBS. | 1:500<br>1:1000 | Abcam; UK |
| Secondary | Goat anti-Rabbit<br>Goat anti-mouse<br>in PBS-T mix | 1:10,000<br>1:10,000 | Licor; USA<br>IRDye® 800CW and<br>680CW, respectively. |

### Western Blotting

For protein extraction, cultured primary pericytes were lysed with cold lysis buffer of radioimmunoprecipitation assay (RIPA) lysing buffer supplemented with 1x protease/phosphatase inhibitor (Cell Signaling Technology; U.S.A). Cells were mechanically agitated for 5 minutes and incubated on ice for a further 30 minutes with occasional agitation. The cell suspension was collected and centrifuged at 4°C for 15 minutes at 10,000g and cell protein extracts were stored at −80°C or brought forward for total protein quantification using Bradford assay (Bio-Rad; USA). Cell protein samples were suspended in 2x Laemmli buffer (Bio-Rad; USA) consisting of 5% 2-mercaptoethanol (ThermoFisher Scientific; USA). 40ug of each sample was denatured at 95°C for 5 minutes and loaded in 4–20% Mini-PROTEAN® TGX™ Precast protein gels (Bio-Rad; SA) with a gel lane including a Dual Xtra standard protein ladder (Bio-Rad; USA). Proteins transferred onto low fluorescence PVDF membranes (ThermoFisher Scientific; USA) running over night at 4°C. Following transfer, PVDF membranes were blocked (5% dried milk powder/TBS-T) for 1 hour at room temperature and incubated at 4°C overnight in primary antibody (Table 2).

Primary antibody mix was removed and PVDF membrane washed 3x times in TBS-T. Secondary antibody mix (Table 2) were incubated on PVDF membrane, light protected for 1 hour. After incubation, PVDF membrane was washed 3x times in TBS-T and visualised using the Odyssey DLx imager (LI-COR Biosciences; USA).

### Statistical analysis

All data are represented as mean±SEM unless stated. Data were acquired and quantified using Microsoft Excel, Image J (https://imagej.nih.gov/ij/) (Schindelin *et al*., 2012; Schneider *et al*., 2012) and Graphpad Prism 9. For *in vitro* calcium assays post AngII readings were determined as a fold change over baseline (T=0) and timeline data were used to provide area under curve analysis (AUC). Immunofluorescent staining of western blotting, images were captured on a Licor Odyssey and analysed Image J software. Protein expression was determined as fold change relative to housekeeping protein. Nociceptive behavioural analysis was performed using a two-way ANOVA with Bonferroni’s multiple comparisons test.

Confocal image processing for data extraction and analysis; Before the analysis of each image, each image was processed within ImageJ v. 2.0.0. software. Images were opened as Z-stack hyperstacks. Hyperstack images were converted into a Z projection to form a 2-dimensional image. Immunofluorescent analysis was performed on ImageJ software including cell counts, K plot value measurements and fluorescent signal for integrated density. To perform a count of structures: the plugin feature of cell count was selected per image and selected. Once selected a recording was made and highlighted on the image. K plot value measurements: This feature was adopted to record the diameter of CD31 labelled vessels, NG2 labelled pericytes and COLIV labelled basement membranes. Before the use of this feature, an image with a known scale bar was used to calibrate the pixels of an image to a known distance. A straight line was drawn cross a structure and the K plot of the straight line was plotted. Gray value peaks representing fluorescent capture within the K plot were measured peak to peak to provide diameter measurements. As part of pericyte characterisation, CD31 labelled vessel diameter measurements were taken. To determine the region of the CD31 labelled vessel region directly beneath the NG2 labelled pericyte, NG2/DAPI expressive pericyte cells positioned on a CD31 labelled vessels were identified. Within CD31 only channels, CD31 labelled vessel regions directly superimposed beneath a NG2 labelled pericyte soma head were measured using the K plot method. This was repeated along the pericyte process 10µm increments in distance away from the NG2 labelled pericyte soma head measurement on the same vessel. This allowed for NG2 labelled pericyte soma and end feet constriction analysis within spinal dorsal tissue regions. To measure integrated density of multiple channels of neuronal cells, the freehand selection tool from ImageJ was selected and drawn around NeuN labelled structures within 1024×1024 pixel images. NeuN labelled neurons were drawn around and added to ROI manager plugin of ImageJ. This allowed for ROI drawn outline to be reapplied on a single channel and analysed to provide exact colocalised readings.

Laser speckle imaging; The moorFLPI-2 instrument was calibrated before using the MOORFLPI_CAL 2PFS calibration block. Imaging the exposed spinal cord through laser speckle microscope were analysed through the moorFLPI-2research software allowing the mean flux (change in blood flow normalised by baseline values) to be determined from ROI selection of spinal cord within the image. Mean flux measurements were taken before topical treatment of drugs as baseline and post −1,-3,-5 and −10 minutes of AngII treatment. Once images were taken the animal was immediately terminated via the S1 procedure of cervical dislocation to end study.

Intravital imaging; Acquired images were captured (.LIF) and exported using Leica LAS software (acquired image precision of either 256 x 256 pixels or 512 x 512 pixels). These were converted into .TIFF or .avi file format. Analysis for measurement of red blood cell velocity (RBC) velocity over time, a region of the vessel lacking bifurcations and overlap of ROIs between differing vessels was chosen. Vessel diameter was recorded by measuring across the segment of identified vessel and a plot profile was produced to evaluate fluorescence intensity. Vessel diameter was determined by measuring between the two fluorescent peaks, which identify vessel wall. Sodium fluorescein was used to measure RBC velocity using kymographs of cell trafficking through the vessel of known radius. Velocity of blood flow was determined by imaging sodium fluorescein filled vessels to track RBC movement along the vessel lumen. Unlabelled RBCs identified by shadows (areas of no fluorescent molecule in the lumen) were used to produce kymographs depicting x axis representing distance (Δd) and y axis represents time (Δ*t*).

## Results

The anti-hypertensive agent, Angiotensin II (AngII), was applied to the spinal cord and blood flow was measured using intravital imaging of the mouse spinal cord. Dorsal horn capillary diameter was determined following WGA and sodium fluorescein capillary perfusion (representative image Fig. 1A). A fluorescence profile plot across the capillary was used to determine fluorescence intensity peaks of WGA and fluorescein was produced and capillary diameter was determined from measuring peak to peak (Fig. 1B). 100nM AngII led to a reduction in dorsal horn capillary diameter when compared to vehicle treatment (Fig. 1C). Additionally, capillary perfusion was measured by determining movement of red blood cells (RBC) through the capillary, with perfused/flowing vessel identified by movement of RBC through the vessel (Fig. 1D), with vessels deemed to have stalled represented by RBCs that did not move (Fig. 1E). Movement of RBC through the capillary was recorded allowing for the measurement of RBC velocity as well as to determine the number of flowing or stalled vessels. AngII led to a reduction of red blood cell movement (Fig. 1F) as well as increased numbers of stalled capillaries (Fig. 1G). 30 minutes following i.t. administration of either vehicle (PBS) or AngII (100nM), the endothelium (CD31) and pericytes (NG2. DAPI) were identified in the dorsal horn (Fig. 2A representative images). This allowed identification of unconstricted and constricted pericytes to be identified (Fig. 2B representative images). Vessel diameter beneath pericytes was measured to categorise pericyte contractility at cell soma and cell process. Vessel diameter beneath pericyte somas (6.02µm±0.11) were larger than vessels beneath the pericyte process (4.72µm±0.1) indicated unconstrictive pericytes (Fig. 2C,D). Constrictive pericytes were categorised when vessel diameters beneath pericyte somas (4.79µm±0.1) were narrower than vessel diameter beneath pericyte process regions (5.84µm±0.12) (Fig. 2E,F). AngII led to an increased number of constricted pericytes in relation to vehicle treated (Fig. 2G), and there were no alterations in the number of pericytes identified in each experimental treatment group. Additionally, there was no significant change in large vessel diameter (Fig. 2H) or count of contracted vessels in either vehicle or AngII experimental groups (Fig. 2I) that were identified as arterioles (pre-capillary vessels SMA positive, Fig. 2J).

**Figure 1.**
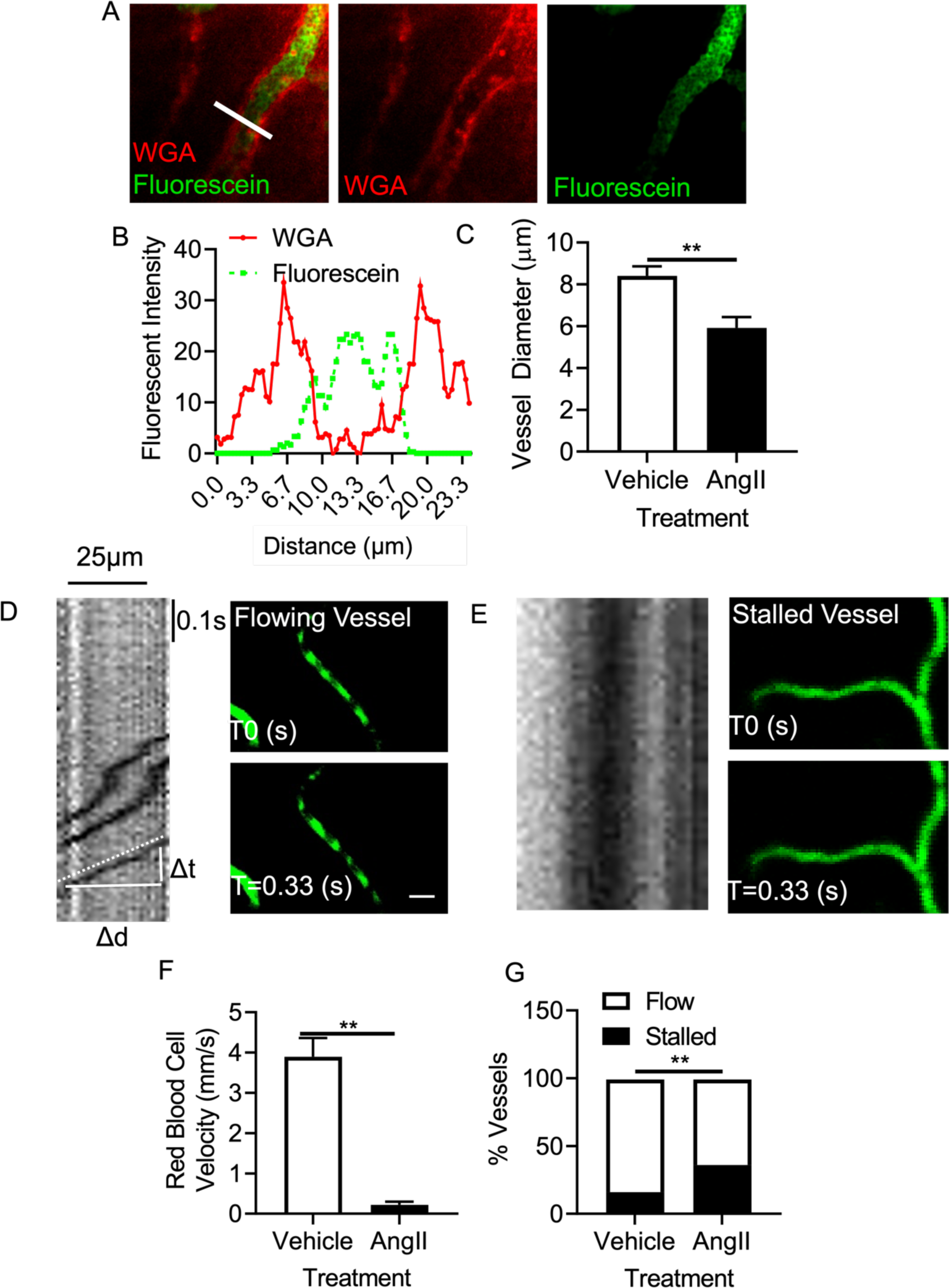
Angiotensin II mediated vasoconstriction in the dorsal horn of the spinal cord. Using intravital imaging via a custom built stereotaxic frame and confocal microscopy, the dorsal horn microvasculature in an adult C57 Bl6 male mouse was imaged to determine levels of capillary perfusion. [A] Microvessels in the dorsal horn were identified following intravenous administration of WGA Alexafluor 555, as well as Sodium Fluorescein (100mg/ml) administered via intraperitoneal injection to enable visualisation of RBCs. [B] Capillary diameter was determined from fluorescent profile plots acquired from a region of interest drawn across the vessel to allow peak to peak measurement of luminal walls. [C] AngII led to a reduction in dorsal capillary diameter versus vehicle control (**p<0.01, Unpaired T Test). RBCs can be visualised a shadows (lack of fluorescein staining) [D & E]. RBC movement were tracked as they move through a [D] perfused vessel (‘Flowing’) and in [E] unperfused (‘Stalled’) vessels where they do not move. Furthermore, [F] Administration of 100nM AngII led to the RBC velocity through microvessels in the dorsal horn decreasing compared to vehicle control (**p<0.01, Unpaired T Test). [G] Furthermore, the number of stalled RBCs was increased in capillaries post AngII treatment when compared to control. (AngII: angiotensin II, RBCs: red blood cells, WGA: Wheat Germ Agglutin)

**Figure 2.**
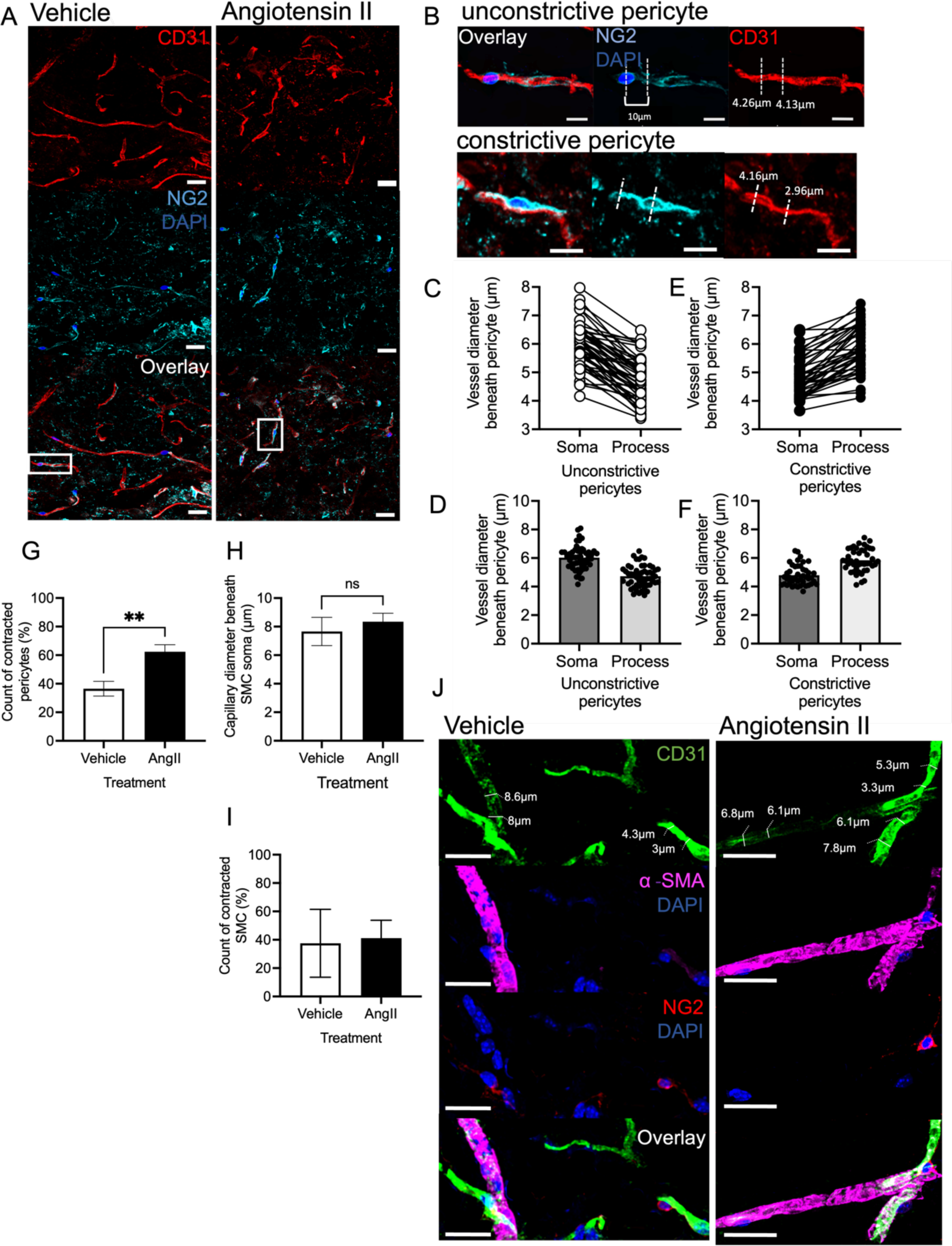
Angiotensin II induced pericyte vasoconstriction in the dorsal horn. Immunofluorescent labelling of transverse PFA-fixed spinal cord sections from either i.t. injected PBS or AngII groups was performed to investigate acute AngII induced-microvasculature changes. [A] Representative images of capillaries in the dorsal horn. Scale bar at 25µm. Pericytes (cyan-NG2, blue-DAPI) were identified by proximal positioning to capillary endothelium (green-CD31). Change in microvessel diameter mediated by pericytes was determined through vessel diameter measurements proximal to the central position of a pericyte soma and compared to the vessel diameter 10µm increments away from the soma (Scale bar 10µm). [B] Pericyte contractility in a region of interest was determined via comparing vessel diameter at the cell soma to along the cell process 10µm increments away. [C&D] Pericytes depicted as unconstrictive showed vessel diameters beneath pericyte somas equal to or thicker than vessel region beneath the pericyte process, whereas [E&F] a contracted pericyte is indicated by the vessel diameter being narrower at the pericyte soma compared to 10μm increments along the pericyte process. [G] Increased number of pericytes were demonstrated to show a reduction in vessel diameter ie vasoconstractility in AngII treatment group compared to the vehicle 30 minutes post-i.t. dosing. To determine pericyte drop off, count of pericytes between treatment groups were normalised to total count of vessels within dorsal spinal window of view (254µm^2^ ROI). No statistical change was apparent between i.t vehicle and AngII treatments. [H] In larger α-SMA positive vessels post-angiotensin II delivery there was no change in vessel diameter compared to vehicle treatment. Furthermore, [I] the percentage of contracted pericytes following AngII treatment in the spinal cord was significantly increased compared to vehicle treatment (**p<0.01). [J] Representative images of vehicle and AngII arterioles and pericytes upon capillaries. A top panel of CD31 staining represents a non-contracted SMC in vehicle treatment group where the position of the SMC soma is proximal to a vessel diameter of 8.8µm. 10µm increments downstream of SMC soma, vessel diameter is 8µm. No statistical difference of contracted SMC was shown between treatment groups. Whereas in AngII treatment group, both a non-contracted SMC (6.1µm vessel diameter proximal to soma, 6.1µm vessel diameter 10µm increments from soma) and a contracted SMC (6.1µm vessel diameter proximal to soma, 7.8µm vessel diameter 10µm increments from soma) are represented. Panel A also represents in vehicle treatment group a non-contracted pericyte within NG2 and DAPI staining (4.3µm vessel diameter proximal to soma, 4.5µm vessel diameter 10µm increments from pericyte soma). Data shown are mean± S.E.M. Unpaired parametric T-test applied. (α-SMA, α-smooth muscle actin, AngII: angiotensin II, CD31: Platelet/endothelial cell adhesion molecule-1, DAPI: 4′,6-diamidino-2-phenylindole, pfa: paraformaldehyde, SMC: smooth muscle cell)

Primary spinal cord isolated pericytes (NG2 labelled Fig. 3A & B) were isolated and calcium handling following AngII treatment was investigated, with isolated pericytes expressing AT1 (Fig. 3C-E). Dose-dependent administration of increasing concentrations (vehicle, 1nM, 30nM and 100nM) of AngII led to increased calcium influx in isolated spinal cord pericytes, presented as increased intracellular calcium fluorescence over-time (Fig. 3F). In addition, AUC analysis of intracellular calcium demonstrated 100nM AngII increased fluorescence intensity compared to vehicle treatment (PBS) (Fig. 3G). Immunofluorescent staining of mouse dorsal horn demonstrated pericytes (NG2 labelled) in close contact to the endothelium (CD31) colocalising with AT1 staining (Fig. 3H).

**Figure 3.**
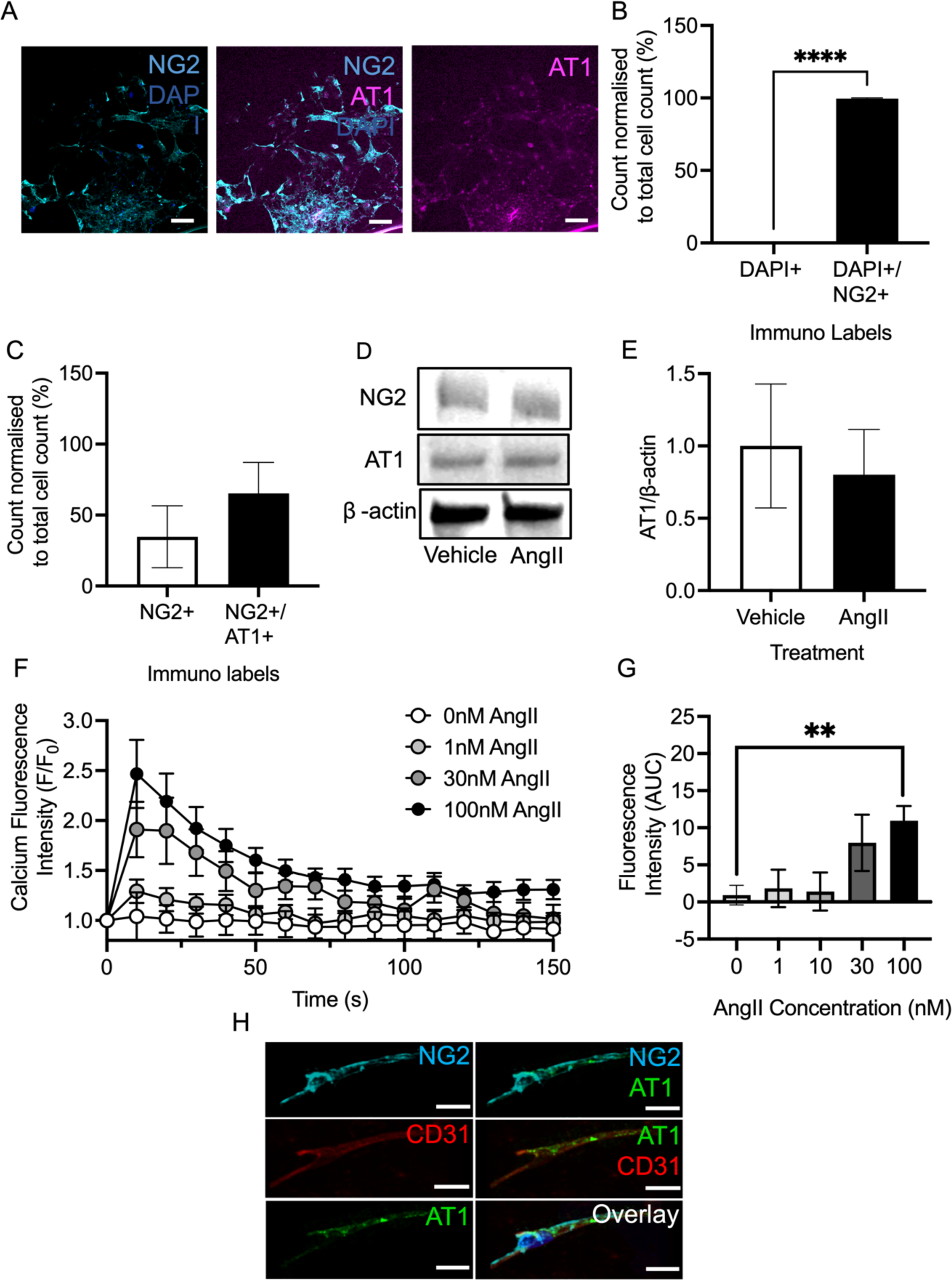
Angiotensin II induced calcium influx spinal microvascular pericytes. [A] *In vitro* spinal pericyte cultures were established and immunofluorescently stained with NG2, DAPI and AT1 (Scale bar at 20µm). [B] Identification of spinal mouse pericyte isolation was provided by percentage count of DAPI staining alone compared to NG2 and DAPI positive stained cells, indicating a 100% count of NG2 and DAPI colocalization. [C] Percentage count of cells expressing AT1 co-stained with NG2 and DAPI in pericyte cultures indicated no statistical difference between NG2 only and NG2 and AT1 positive cells cultured in normal media. Western blotting further validated pericyte cell cultures with 50µg load protein lysate expressing [D&E] NG2 (240kDa) alongside expression of AT1 (52kDa) protein band. Calcium intracellular movement is a key event during contractile modulation in AngII-induced pericyte constriction. Treatment of AngII at concentrations of 0,1, 30 and 100nM (n=6-9), primary spinal pericyte cell cultures indicated a dose-dependent effect of increased intracellular calcium activity [F] over time and represented as [G] AUC from calcium fluorescent intensity change (*p<0.05 Mann-Whitney test). [H] Immunofluorescent labelling of transverse PFA-fixed spinal cord tissue sections indicated pericytes expression of AT1 receptor via colocalisation of AT1 and NG2 staining (Scale bar at 10µm). (AngII: angiotensin II, AT1: angiotensin II type 1, DAPI: 4′,6-diamidino-2-phenylindole, PFA: paraformaldehyde)

To investigate the implication of AngII-induced pericyte contractility upon nociceptive processing, nociceptive behavioural tests were performed. Mechanical and thermal stimuli evoked-paw withdrawals were investigated 1 hour and 24 hours following intrathecal administration of either AngII or vehicle. Mechanical withdrawal thresholds (Fig.4A) and heat withdrawal latency (Fig.4B) was significantly reduced at 1 hour post-AngII administration compared to vehicle groups, with mechanical withdrawal thresholds remaining reduced for up to 24 hours post i.t. AngII treatment.

**Figure 4.**
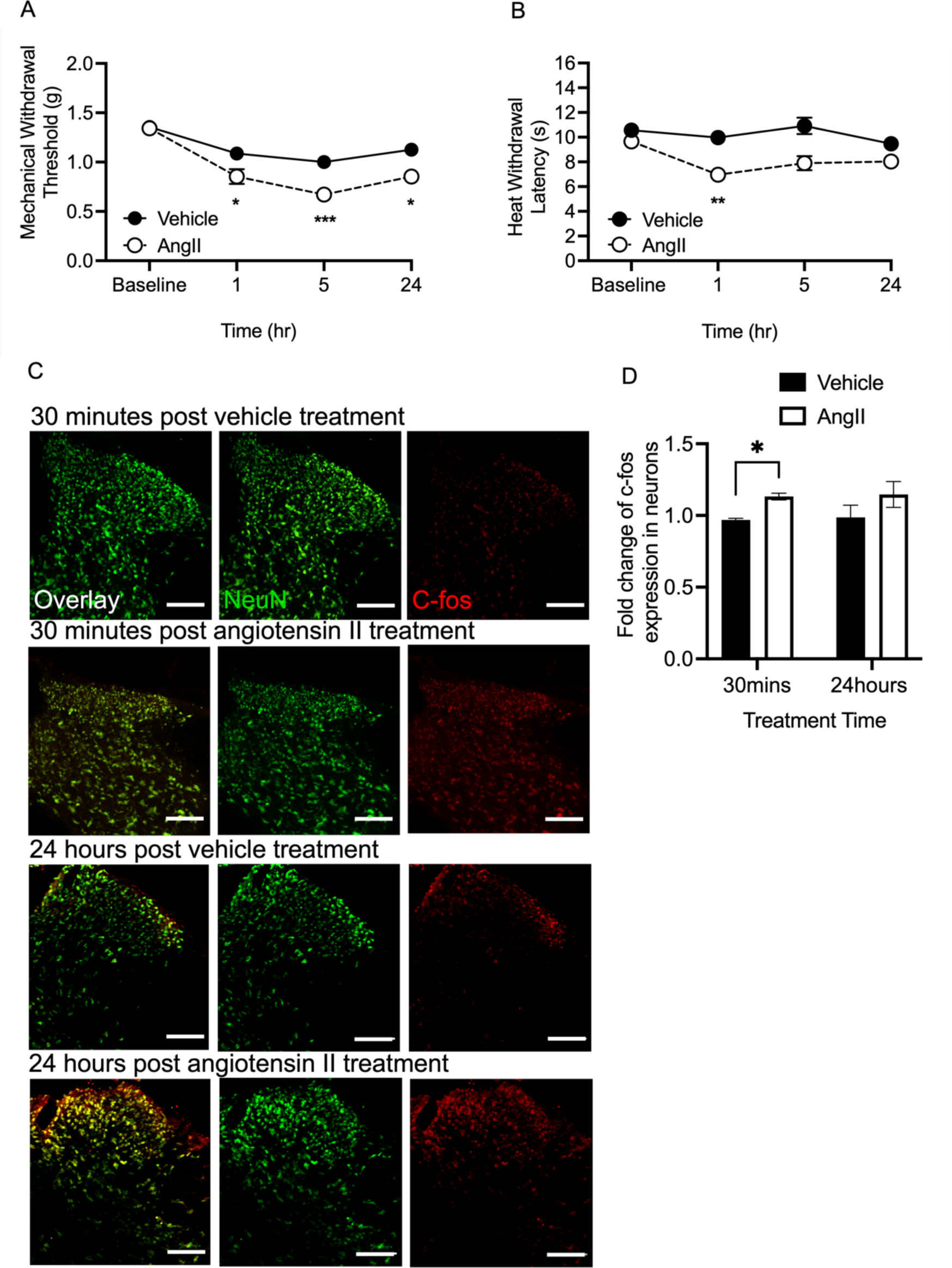
Angiotensin II induces dorsal horn neuron activation and pain hypersensitivity. Behavioural tests of [A] Von Frey filament and [B] Hargreaves were performed on intrathecally dosed C57/Bl6 male and female mice of either i.t. vehicle (PBS, n=30) or AngII (100nM, n=21) to determine the effects of spinal dorsal vascular change on nociceptive signalling. Both mechanical (*p<0.05) and thermal-evoked (**p<0.01) paw withdrawal thresholds significantly reduced below vehicle treatment thresholds post-1hr of i.t. AngII administration. Hypersensitivity was detected at 5hrs (***p<0.001) and 24hrs (*p<0.05;Two-way ANOVA t-test with post-hoc Bonferroni test). [C] Immunofluorescent labelling of transverse PFA-fixed spinal cord tissue was performed. Tissue harvested at 30 minutes (n=6) and 24 hours (n=6) post-i.t. injected with vehicle (n=3) and AngII (n=3) were labelled with NeuN (green) and c-fos (red). C-fos positive NeuN neurons were used to determine drug-induced activation of dorsal horn neurons (Scale bar at 100µm). [D] C-fos positive NeuN counts in the dorsal horn were recorded for each experimental group and normalised to total NeuN count. Significant increase in fold change of c-fos was detected 30 minutes post-AngII dosing (*p<0.05) compared to 30 minutes-post vehicle dosing. In each time point group, counts in vehicle groups were used to calculate fold change across vehicle and AngII counts (*p<0.05. Multiple unpaired parametric t-test, n=3 per group). (AngII: angiotensin II, PBS: phosphate buffer saline, PFA: paraformaldehyde)

**Figure 5.**
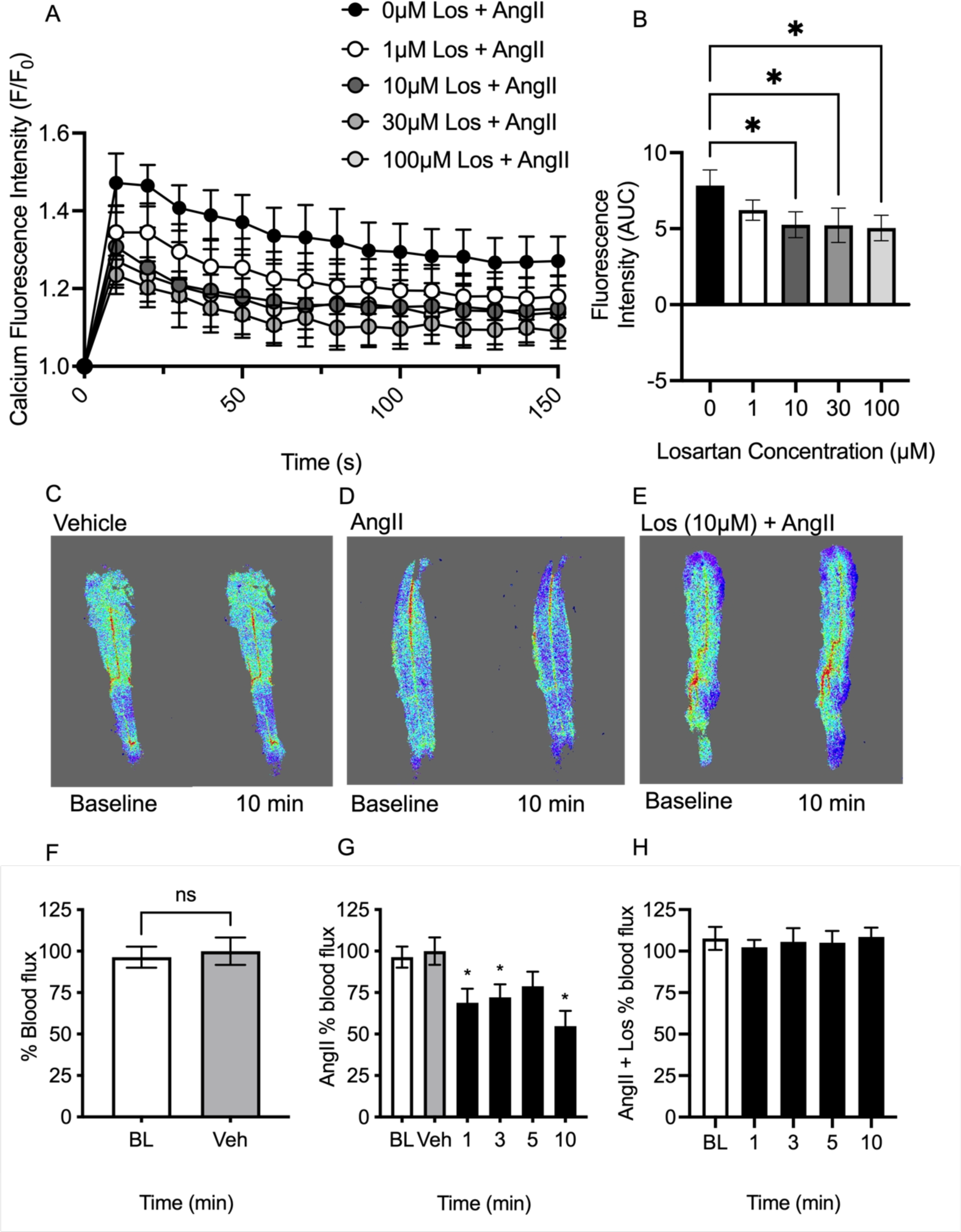
Losartan attenuates angiotensin II-induced activation of pericytes and angiotensin II-induced spinal blood flux reduction. To investigate the effects of angiotensin II type 1 receptor inhibition an AT1 inhibitor, losartan, was introduced alongside AngII (100nM) in cultured primary spinal cord pericytes at varying concentrations of 0, 1, 10, 30 and 100μM. [A] Overtime, a dose dependent effect of reduced calcium fluorescent intensity was detected as losartan concentrations increased when co-administrated with AngII representing reduction in pericyte activation. [B] AUC indicated significant change in calcium fluorescent intensity at 10, 30, and 100µM losartan treatment compared to AngII dosed without losartan (*p<0.05, One-way ANOVA with post hoc Bonferroni test applied, N=4). Blood flux was investigated using Laser Speckle contrast microscopy in anaesthetised (2% isofluorane) male C57/B6J mice. Laminectomy at lumbar region was performed on anaesthetised mice and blood flow was recorded pre-dosing as a baseline and at 10 minutes post-dosing with [C] vehicle (PBS, n=10), [D] AngII (100nM, n=10) or [E] los (10µM, n=10) + AngII (100nM). Percentage blood flux were calculated at 1, 3, 5 and 10 minutes post drug dosing. [F] Vehicle treatment blood flux values were recorded and plotted at 10 minutes post-dosing with no change post vehicle treatment. [G] AngII treatment post 1 (*p<0.05), 3 (*p<0.05), and 10 minutes (*p<0.05) indicated a significant reduced from baseline. [H] These changes in blood flux were alleviated over 10 minutes of recordings in animals dosed with losartan alongside AngII (*p<0.05. Unpaired parametric t-test). (AngII: angiotensin II, AT1: angiotensin II type 1, BL: baseline, los: losartan, Veh: vehicle)

The fold change of c-fos positive dorsal horn neurons (NeuN co labelled) in the dorsal horn (Fig.4C representative images) was increased 30 minutes post-AngII treatment compared vehicle treatment (Fig.4D). Post-24 hours, the number of c-fos positive neurons was not different between vehicle and AngII treatment groups.

In isolated spinal cord pericytes, AngII-induced intracellular calcium was determined to be AT1 dependent. Isolated pericytes were treated with 100nM AngII to induce calcium influx with losartan (AT1 receptor inhibitor). Losartan treatment in a concentration dependent manner decreased AngII induced intracellular calcium intensity (Fig. 4A & B). Subsequently, AngII influence on blood flow was explored using Laser Speckle Image Contrast microscopy of the dorsal horn. Measurements of blood flux were performed in vehicle, AngII and AngII plus losartan (Fig. 4C, D, E representative traces). Vehicle treatment showed no significant change in blood flux post-dosing (Fig. 4C,F). However, a significant reduction in blood flux was present at 1, 3 and 10 minutes post-AngII dosing compared to baseline values (Fig. 4D, G). Co-administration of losartan (10µM) with AngII prevented AngII induced suppression in blood flow (Fig. 4E, H)

Consequently, AngII modulation of nociceptive behaviour was determined to be AT1 dependent, following intraperitoneal administration of AT1 (Losartan) or AT2 (PD12331) inhibitors. Losartan (i.p., 20mg/kg) was administed via intraperitoneal injection alongside AngII (i.t., 100nM) versus i.t. AngII dosed alone. There was no difference in nociceptive behavioural withdrawals induced by i.p. Losartan alongside i.t. AngII in mechanical evoked paw withdrawal thresholds compared to AngII group (Fig. 6A). Whereas i.p. Losartan delivered alongside i.t. AngII induced an increase in heat-evoked withdrawal latency in animals compared to animals dosed with i.t. AngII alone (Fig. 6B). When exploring the effect of AT2 antagonism in the presence of AngII upon nociceptive behaviours, administration of PD123319 (i.p. 10mg/kg) alongside i.t. AngII led to no change in the i.t. AngII induced mechanical (Fig. 6C) and heat (Fig, 6D) hypersensitivity. In addition, AngII induced pain behavioural hypersensitivity to mechanical and heat stimuli in both male and female mice. Furthermore, AngII induced pain was AT1 dependent at the level of the dorsal horn in both male and female mice. Female mice intrathecally dosed with AngII showed a significantly reduced withdrawal threshold to mechanical evoked stimuli (Fig. 7A) and heat evoked withdrawal latency (Fig. 7B) compared to vehicle treatment groups. Following this, female mice dosed with i.t. Losartan alongside i.t. AngII indicated a significant increase in mechanical withdrawal threshold (Fig. 7C) and heat withdrawal latency (Fig. 7D) compared to AngII alone. When comparing AngII mediated alterations in pain behaviours in male mice, mechanical withdrawal threshold (Fig. 7E) and heat evoked paw withdrawal latency (Fig. 7F) were significantly reduced in mice dosed with i.t. AngII compared to the vehicle treatment group. The inclusion of i.t. losartan dosed alongside i.t. AngII induced a significant increase in mechanical withdrawal threshold (Fig. 7G) and heat evoked withdrawal latency (Fig. 7H) compared to male mice dosed with i.t. AngII alone.

**Figure 6.**
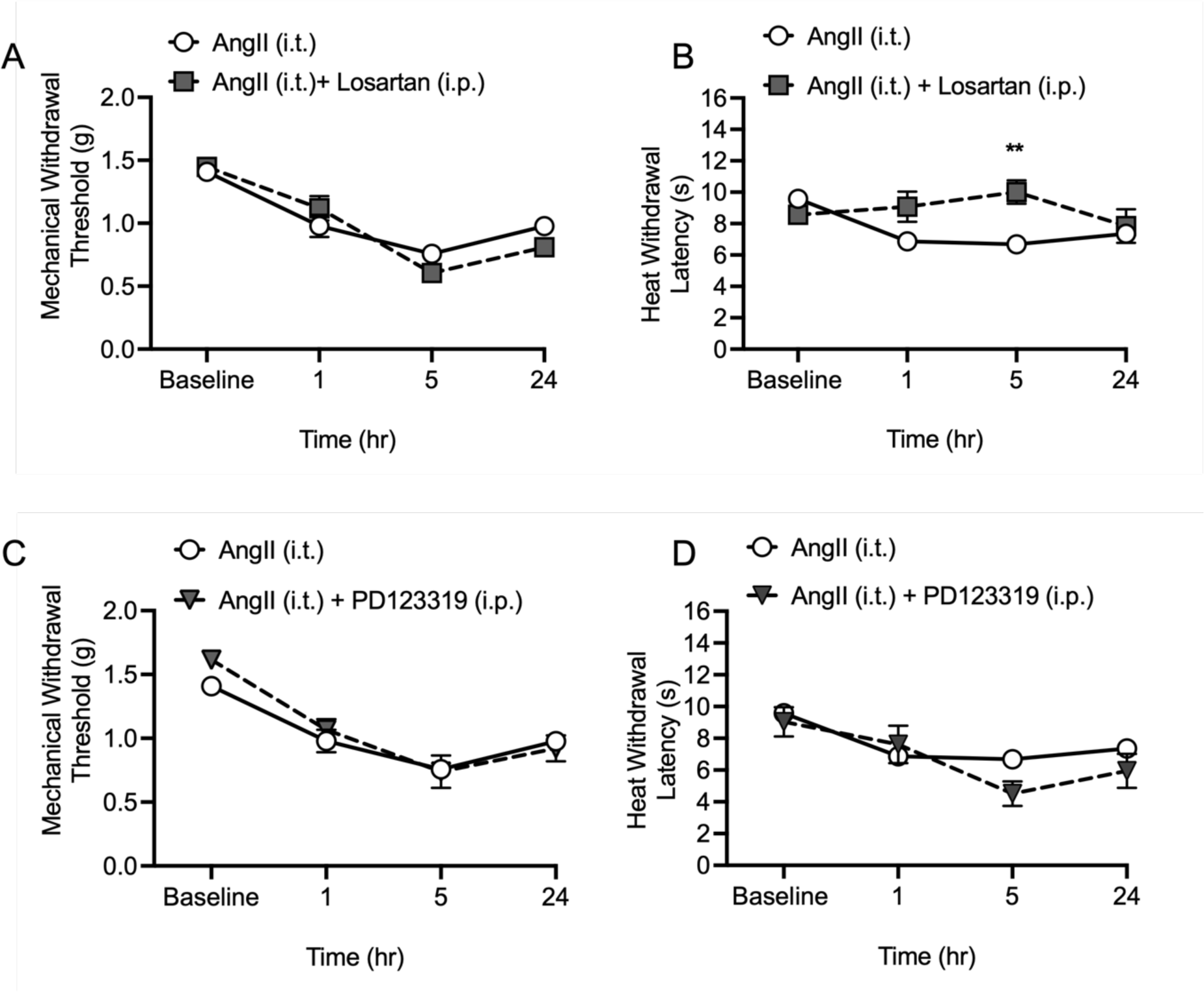
Angiotensin II-induced nociceptive hypersensitivity is AT1 dependent. Prior to behavioural testing, i.p dosing of losartan (20mg/kg, n=6) or PD123319 (10mg/kg, n=3) was administered in anaesthetised mice alongside i.t. AngII (100nM) to determine if hypersensitivity is elicited via angiotensin II type 1 or 2 receptor activation. [A] Mechanical withdrawal threshold from Von Frey filament test in mice dosed with i.t. AngII and i.p. Losartan showed no differences between i.t. AngII alone over 24 hours. [B] However, AngII induced thermal hypersensitivity was significantly alleviated by i.p losartan compared to i.t. AngII dosing alone post-5 hours (**p<0.01). I.p. PD123319 dosed alongside i.t. AngII did not significantly alleviate AngII induced hypersensitivity from Von Frey filament mechanical-stimuli testing [C] or Hargreaves thermal-stimuli testing [D] over 24 hours. Data shown here are mean ± S.E.M. Two-way ANOVA t-test with post-hoc Bonferroni test applied. (AngII: angiotensin II, i.p.: intraperitoneal, i.t.: intrathecal)

**Figure 7.**
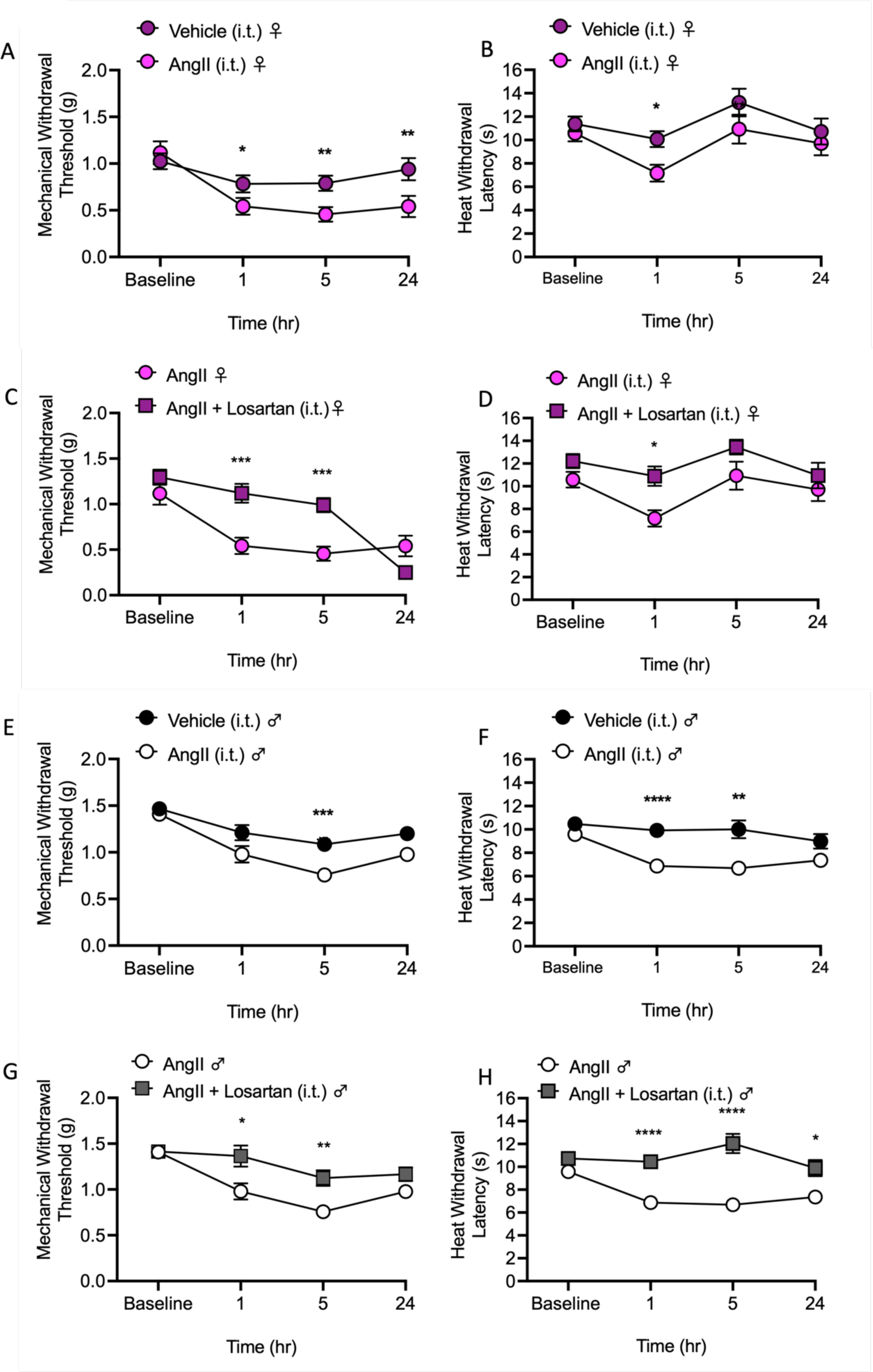
Angiotensin II-induced hypersensitivity in male and female mice model is AT1 dependent at the level of the spinal cord. To determine if sexual dysmorphism is present in AngII-induced nociceptive hypersensitivity the withdrawal thresholds of female C57/Bl6 to mechanical and thermal stimuli were compared against male nociceptive behavioural responses. Moreover, analgesic effects mediated by angiotensin II type 1 inhibition via i.t. Losartan coadministrated with i.t. AngII was investigated. Female mice demonstrated a reduction in withdrawal threshold in response to [A] mechanical stimuli and [B] withdrawal latency to heat stimuli from i.t. vehicle treatment responses (PBS, n=6) 1hour post i.t. AngII delivery (100nM, n=6, *p<0.05). [C] Mechanical induced hypersensitivity was alleviated between 1 (***p<0.001) and 5 hours (***p<0.001) post i.t. AngII and Losartan dosing (10µM, n=3) compared to i.t. AngII delivered alone. In addition, withdrawal latency to heat was increased to female mice post-i.t. AngII and Losartan dosing (*p<0.05) compared to the female mice dosing with i.t. AngII alone. [E] Male mice dosed with i.t. AngII (n=15) indicated a reduction in withdrawal threshold to mechanical stimuli compared to i.t. vehicle treatment groups (***p<0.001) n=15) at 5 hours post-dose. [F] Male mice demonstrated a reduction in heat withdrawal latency at 1hour (****p<0.0001) and 5hours (**p<0.01) post i.t. AngII dosing compared to i.t. vehicle treatment group. [G] Intrathecal administration of losartan alongside AngII in male mice, alleviated the reduction in withdrawal threshold to mechanical stimuli at 1hour (*p<0.05) and 5 hours (**p<0.01) post-dosing compared to i.t. AngII dosing alone. [H] The alleviation of reduced withdrawal latency to heat was detected over 1 (****p<0.0001), 5 (****p<0.0001) and 24 (*p<0.05) hours post i.t. AngII/Losartan dosing compared to i.t. AngII dosing alone (Two-way ANOVA with post-hoc Sidak testing). (AngII: angiotensin II, PBS: phosphate buffer saline, i.p.: intraperitoneal, i.t.: intrathecal)

Basement membrane (BM, COLIV) and endothelium (CD31) were labelled and evaluated to investigate the integrity of the dorsal horn vasculature following intrathecal administration of vehicle, AngII and AngII+losartan treatment (Fig. 8A). Post-24 hours of dosing, i.t. AngII dosed alone indicated a pronounced endothelial regression and fragmentation. There was a reduction in size of CD31 positive vessel fragments (in relation to basement membrane sleeves) in the dorsal horn compared to vehicle (PBS) and i.t. AngII with Losartan treatment group (Fig. 8B). In addition, there was a significant reduction in the endothelium diameter versus diameter of BM in the dorsal horn of the i.t. AngII dosed group (Fig. 8C). Vehicle and i.t. AngII with Losartan treatment groups showed no significant difference between vessel and BM diameter measurements.

**Figure 8.**
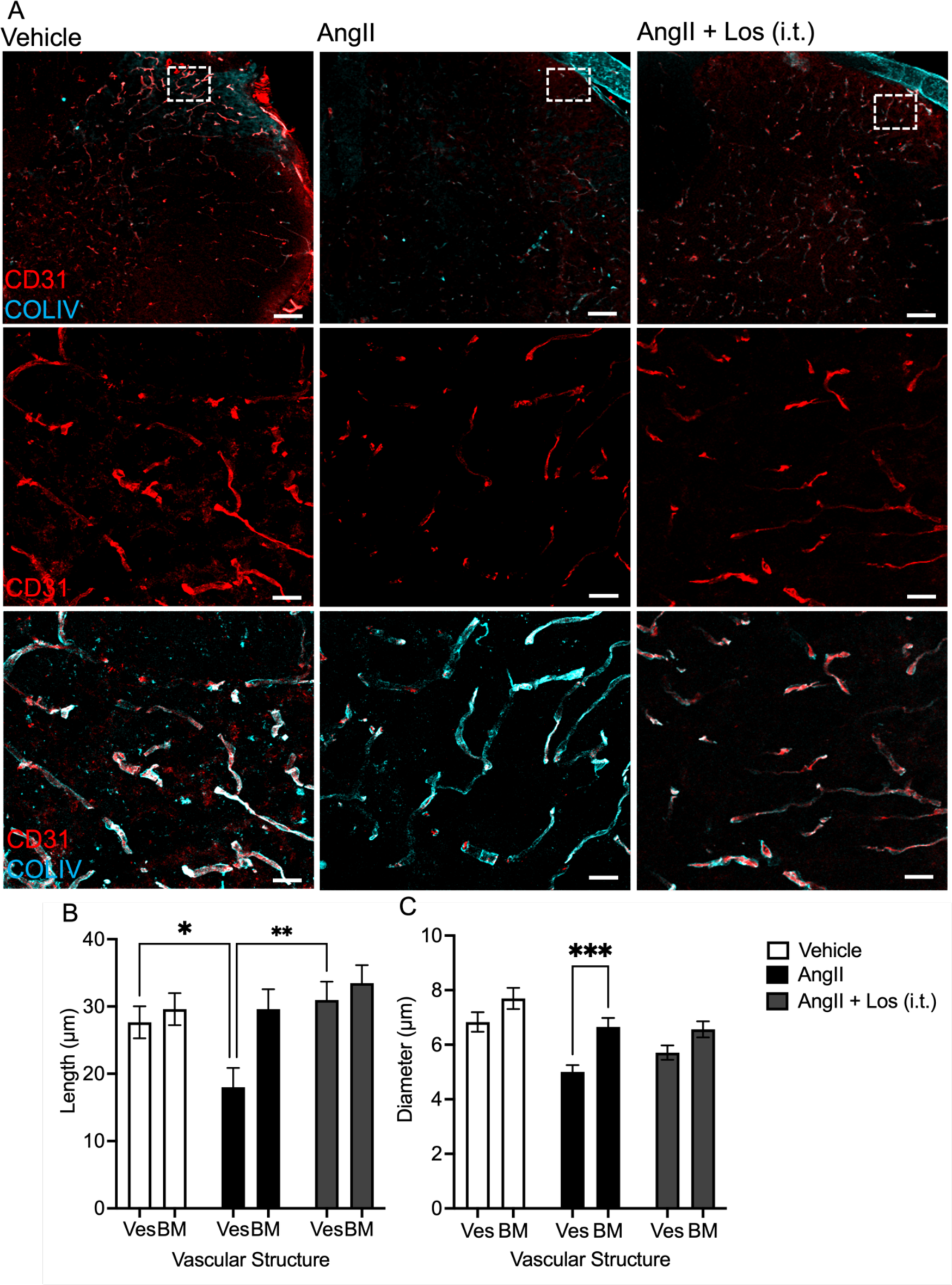
Angiotensin II-induced endothelium deterioration in the dorsal horn is AT1 mediated. [A] Representative examples of immunofluorescent staining was used to determine the integrity of PFA-fixed spinal lumbar dorsal horn endothelium (CD31, red) and basement membrane sleeves (COLIV, cyan) in animals dosed with vehicle, i.t. AngII (100nM, n=5) or i.t. AngII combined with los (10µM, n=3) (Scale bar at 100µM and 25µM) [B] AngII led to a deterioration in the dorsal horn endothelium (CD31 stained) demonstrated as a reduction in endothelial fragment size versus basement membrane sleeve when compared to vehicle treated and AngII + los treated groups. [C] To support the impact of AngII on vessel integrity, endothelium diameters were measured and compared to BM diameter. Ves diameter following i.t. AngII treatment showed a significant reduction compared to BM diameter (***p<0.001; One-way ANOVA with post hoc Bonferroni). (AngII: angiotensin II, BM: basement membrane, CD31: Platelet/endothelial cell adhesion molecule-1, COLIV: collagen IV, los: losartan, pfa: paraformaldehyde, Ves: vessel)

## Discussion

Vascular disturbances have proven to be fundamental in modulating nociceptive processing and underlie the development of chronic pain states(Younis *et al*., 2022). However, investigations into the control of vascular function, in particular vascular tone have not been extensively explored in the context of nociception and onset of chronic pain states. The contribution of mural cell, involvement in particular pericytes, have been widely implicated in neurodegenerative disorders. Here we demonstrate an angiotensin II induced pericyte mediated modulation of spinal cord blood flow, via AT1 dependent vasoconstriction. Furthermore, this AngII-AT1 driven vasocontractility in the dorsal horn initiates pain hypersensitivity.

### Pericyte dependent mediation of pain

There is an increasing recognition that the blood spinal cord barrier plays an important role in controlling nociception as well as instigating the transition to chronic pain states. Previous investigations in particular focussing upon traumatic nerve injury and inflammatory induced pain states, implicate enhanced vascular permeability as a mediator in the onset of chronic pain(Beggs *et al*., 2010). This is typified by the loss of tight junctional proteins and enhanced solute permeability (increased penetration of fluorescent tracers for example), factors that outline the molecular mechanisms that enables inflammatory cell infiltration and pro-inflammatory processes into neural tissues responsible for nociceptive processing (Montague-Cardoso *et al*., 2020; Montague-Cardoso & Malcangio, 2021). In contrast, our work has presented that damage and cessation of blood flow in the spinal cord vasculature also initiates chronic pain states(Ved *et al*., 2018*a*). This is further emphasised by the induction of hypoxia in the microenvironment of the dorsal horn neurons driving chronic pain states(Da Vitoria Lobo *et al*., 2022). The spinal cord vasculature, which comprises of largely of endothelial cells (EC), degenerates either through endothelium apoptosis or regression, and is implicated in the onset of numerous neurological diseases. Our work presented to date highlights that if the endothelium degenerates exclusively, pain manifests(Ved *et al*., 2018*a*; Da Vitoria Lobo *et al*., 2022). However, it remains undefined the cellular and/or molecular events that underpin the diminishing health of the dorsal horn endothelium.

Dorsal horn sensory neurons possess this maladaptive plasticity to enable pain systems to respond to trauma or lesion, such as presented by ourselves that the dorsal horn endothelium has a contributing role in modulating pain perception(Beazley-Long *et al*., 2018; Ved *et al*., 2018*a*). This vascular degeneration is evident in rodents models of diabetic neuropathic pain with a reduction in spinal cord endothelium. Furthermore, vascular integrity is largely mediated by the Vascular Endothelial Growth Factor (VEGF) family of proteins. VEGF signalling via VEGF receptor 2 (VEGFR2) in part through applied mechanical shear stress as a consequence of blood vessel perfusion. Cessation or reductions in flow drives vascular regression typified by reductions in vascular endothelial growth factor (VEGF) A-VEGF receptor 2 signalling, a system we have utilised using a mouse transgenic model to ablate endothelial VEGFR2 signalling through an endothelial specific VEGFR2KO mouse(Beazley-Long *et al*., 2018; Ved *et al*., 2018*a*). This transgenic VEGFR2KO mouse presents pain hypersensitivity. How blood flow can be altered in the capillary can be elicited by varying means, but narrowing of the capillary luminal diameter alters the haemostatic properties passing over the endothelium.

In line with this, the vasculature is highly adaptable, with the ability to respond to a number of stimuli. Under normal physiological pressures the endothelium is maintained via tissue perfusion in a quiescent state. However, there are extensive investigations that depict alterations in mechanical pressures through fluctuations in flow through the capillary lumen, drives alterations in the endothelial activity and health. It can undergo extensive vascular expansion and remodelling during development and pathological conditions. However, vascular regression (or pruning) typified by endothelium degeneration curtails such expansion, has also been established. This occurs in areas of constricted capillary networks and perturbed blood flow(Korn & Augustin, 2015). Typically, a quiescent endothelium relies upon fluid flow induced mechanical shear stress to maintain vascular architecture, with the physical pressure from blood flow driving angiogenic processes such as endothelial cell migration and/or proliferation. However, reduced blood flow due to occlusion or narrowing (i.e. constriction or blockage) of microvessels induces degeneration of neighbouring endothelial cells to this occlusion; initiating vascular pruning. It is widely presented that the auxiliary supportive cell types that comprises of mural cells (includes smooth muscle cells and pericytes), have an important role in supporting the function and integrity of the endothelium. However, under neuropathological conditions, dysfunction in this coordinated communication is implicated in age and metabolic disturbances in cerebrovascular disease (Taylor *et al*., 2015; Kisler *et al*., 2017). In particular, an extensive body of work now presents that pericyte dysfunction or death, is a characteristic of neuropathology (Cai & Boulton, 2002; Kisler *et al*., 2017; Diéguez-Hurtado *et al*., 2019; Nikolakopoulou *et al*., 2019). The vascular system is heavily implicated to be involved with the onset of numerous neuropathologies, and here we present that an angiotensin II driven contractility of pericytes and subsequent narrowing of spinal capillaries initiates mechanical and heat hypersensitivity mirroring endothelium degenerative mouse models that presented pain hypersensitivity.

### AT1 dependent modulation of spinal cord pericyte function and vasomodulation

Our data presented here demonstrates a pronounced AngII induced pericyte vasocontractility that is implicated in the onset of pain hypersensitivity. In addition, we present that this AngII induced activation of pericytic contractility is AT1 dependent. AngII is widely reported to influence capillary function and tone, with particular this recognised through therapeutic targeting of RAS mediated signalling acting as a primary therapeutic target for treatment of hypertension. Patients with hypertension are increasingly susceptible to neurodegenerative disorders, and in particular chronic pain states (Ponirakis *et al*., 2021) implicating AngII as a causative factor in the onset of chronic pain states. This is supported by the developing notion that pericytes in the nervous system are responsive to RAS signalling and in particular express AT1(Hirunpattarasilp *et al*., 2023). In relation to the spinal cord and nociceptive processing, constituents of the RAS system are expressed extensively in the spinal cord including AngII, ACE and AT1 and in particular associated in chronic pain states(Ogata *et al*., 2016). However, specific cell labelling was not defined in this study, although functionally RAS is involved with pain modulation. Recent evidence highlights AngII dependent modulation of the CNS pericyte-vascular function, with cerebral capillary constriction driven by SARS COV 2 receptor binding domain inducing pericyte contractility in an AT1 dependent manner(Hirunpattarasilp *et al*., 2023). This data indicates a significantly prolonged constriction of pericytes upon AngII treatment, blocked with losartan, supporting the role of AT1 presence and role of pericyte to influence constricted capillary tone. This extended period of contractility has been previously associated with calcium handling, with shown to induce increases in intracellular calcium intracellular in numerous tissue types including podocyte(Edwards & Pallone, 2008; Burdyga & Borysova, 2014; Binz-Lotter *et al*., 2020), with a pronounced increase in intracellular calcium associated with elevated frequency of pericyte contractility(Hirunpattarasilp *et al*., 2022). Additionally, in neurodegenerative states, pericytes can enter a state of ‘rigor’ to induce a permanent state of capillary constrict post-pericyte death (Yemisci *et al*., 2009; Hall *et al*., 2014*b*; Heyba *et al*., 2019), a process dependent upon pericyte intracellular calcium handling(Korte *et al*., 2022). This implies AngII induced pericyte activation of AT1 is fundamental in the regulation of spinal cord capillary tone. The clear indication that losartan protects against AngII induced capillary damage has been demonstrated previously with the AT1 inhibitor, candesartan, preventing AngII induced capillary length and diameter reduction in the cerebral cortex (Lapi *et al*., 2021). Furthermore, prevention of retinal capillary loss has also been observed following losartan treatment (Yang *et al*., 2010; Yu *et al*., 2011; Wu *et al*., 2020) This demonstrates AT1 inhibition could be a putative novel target for analgesia development.

Here we present AngII induced vascular remodelling and modulation of central nervous system processing of nociceptive information in a AT1 dependent manner. However, it must be acknowledged that there have been differing studies that have implicated RAS to be targeted against nociceptor sensitisation, with a focus upon AT2 signalling driving chronic pain(Shepherd *et al*., 2018*c*, 2018*d*). AT1 and AT2 are expressed on dorsal root ganglia sensory neurons, in particularly nociceptors and regulate AngII induced sensory neuritogenesis (Benitez *et al*., 2017, 2020). This is in contrast to our data presented, whereby AngII administration via intrathecal delivery drives chronic pain states. This can be explained by differing approaches to the application of AngII *in vivo*, with much lower concentrations utilised by Shepherd and colleagues during *in vitro* application (Shepherd *et al*., 2018*c*, 2018*d*) and as by us in cellular and *in vivo* models.

### Conclusion

AngII induced alterations in the spinal cord capillary network through changes in blood flow and loss of capillary integrity precedes pain hypersensitivity. Furthermore, AngII exerts activity through AT1 activation expressed upon pericytes. This study depicts AngII as a mediator of chronic pain and acts as a potential therapeutic target in the treatment of chronic pain.

## Acknowledgements

This work was supported by the European Foundation for the Study of Diabetes Microvascular Programme supported by Novartis to RPH (Nov 2015_2 to RPH), the EFSD/Boehringer Ingelheim European Research Programme in Microvascular Complications of Diabetes (BI18_5 to RPH), the Rosetree Trust (A1360 to RPH) and Nottingham Trent University. All authors performed the experimental work and contributed to the conception or design of the work in addition to acquisition, analysis or interpretation of data for the work. All authors drafted the article or revised it critically for important intellectual content. All authors approved the final version of the manuscript. All authors approved the final version of the manuscript, agree to be accountable for all aspects of the work in ensuring that questions related to the accuracy or integrity of any part of the work are appropriately investigated and resolved; and all persons designated as authors qualify for authorship, and all those who qualify for authorship are listed. The authors would like to thank Graham Hickman of the Imaging Suite at Nottingham Trent University for their support and assistance in this work.

